# Defining the pre-symbiotic transcriptional landscape of rice roots

**DOI:** 10.1101/2025.09.25.678570

**Authors:** Gabriel Ferreras Garrucho, Uta Paszkowski, Chai Hao Chiu

**Author notes:** Corresponding authors: Uta Paszkowski, Chai Hao Chiu **Email:**. shared last author (Uta Paszkowski, Chai Hao Chiu). **Author Contributions:** G.F.G., U.P. and C.H.C. designed the study and obtained funding. C.H.C. isolated and characterised the collection of rice mutants. G.F.G. and C.H.C. performed the experiments. G.F.G. analysed the data and generated figures. G.F.G, C.H.C. and U.P. wrote the paper. **Competing Interest Statement:** The authors declare no competing interests.

## Abstract

Plants interact with a plethora of organisms in the rhizosphere, with outcomes that range from detrimental to beneficial. Arbuscular mycorrhizal (AM) symbiosis is the most ubiquitous beneficial plant interaction in terrestrial ecosystems and involves soil borne fungi of the Glomeromycotina. It is believed that plants detect diagnostic signals for the discrimination between beneficial arbuscular mycorrhizal (AM) fungi and parasitic fungi during the pre-symbiotic molecular crosstalk. Here, we investigated the transcriptome of rice roots upon exposure to the complete cocktail of fungal exudates from either beneficial *Rhizophagus irregularis* or pathogenic *Magnaporthe oryzae*. We report that regardless of the exudate donor species, the transcriptional response lacked diagnostic differences. Instead, the profiles were marked by the common suppression of symbiosis signalling components, accompanied by the induction of a generic stress response (GSR) and defense-related signature, which was retained in a suite of symbiosis signalling mutants impaired at different stages of symbiosis development. However, upon permitting physical engagement with AM fungi, a striking reversion in the transcriptional responses occurred marked by the simultaneous relaxation of symbiosis signalling suppression and down-regulation of defense-related and GSR markers, overall comparable between wild-type and mutants. Our data therefore reveal that rather than specific recognition in the rhizosphere, a sequence of signals orchestrates stress, immunity and symbiosis, pivoting towards symbiosis potentially at the stage of plant-fungal contact formation.

**Significance Statement:** Most plant roots engage in symbiosis with beneficial mycorrhizal fungi. Before physical contact, early signal exchanges involve diffusible molecules. The bouquet of chemicals released by beneficial *versus* detrimental fungi has been proposed to convey specificity for signal transduction, transcriptional output and responses. Here we characterised the transcriptomes of rice roots to exudates from beneficial and pathogenic fungi, and found that they lack specific signatures but elicit a generic stress and immune response and suppression of symbiosis signalling. This profile was reversed, along with activation of symbiotic transcriptional responses, only when AM fungi were allowed to establish physical contact. This reveals the need for additional, unknown, layers of signal exchanges for plants to commit to symbiosis.

## Introduction

In both natural and agricultural soil environments, plants are exposed to and interact with a myriad of microbial organisms, with diverse outcomes that range from mutualism to parasitism (1). Beneficial arbuscular mycorrhizal (AM) fungi associate with approximately 85% of terrestrial plant species, delivering inorganic minerals, mainly phosphorus and nitrogen, in exchange for photosynthesised carbon, primarily sugars and lipids (2, 3). However, plants are equally exposed to pathogenic fungi and defend themselves (4). The prevailing view has been that diffusible chitinaceous rhizosphere signals differ between fungi and specify the early activation of symbiosis or immunity signalling in the plant (1, 5, 6). Symbiosis signalling involves the activation of the Common Symbiosis Signaling Pathway (CSSP), so called because of its equal requirement in legumes for the interaction with either AM fungi or nitrogen-fixing rhizobia (7, 8). In contrast, Pathogen Associated Molecular Patterns (PAMPs) such as chitin induce Pattern-Triggered Immunity (PTI)(4, 5, 9). The use of established assays for symbiosis or PTI led to the notion that short-chain chito-oligomers (CO3-5) and lipo-chitooligosaccharides (LCOs) are symbiotic signals and that longer-chain oligomers (CO6-8) are elicitors of PTI (9–14). Perception of chitinaceous molecules involves Lysin Motif Receptor-Like Kinases (LysM-RLKs), of which the functions of Chitin Elicitor Receptor Kinase 1 (CERK1), CERK2, Nod Factor Receptor 5 (NFR5) have been characterised in rice in the context of Microbe Associated Molecular Patterns (MAMP) signalling (15–22). In recent years, it has however become increasingly clear that pathogenic and symbiotic fungi release similar sets of chitinaceous signals, including short and long COs as well as LCOs (5, 23). Moreover, these chitin signals are recognised by overlapping sets of plant receptors, and plants launch similar responses to all of them (9, 24, 25). This calls into question their contribution to the discrete pathogenic and symbiotic outcomes. Conceivably, discrimination between beneficial and pathogenic fungi might be achieved via detection of other fungal exudate constituents during rhizosphere signalling, for instance by recognising proteinaceous effectors that elicit differential responses in the plant (26–30).

To determine whether the plant indeed launches a specific response when exposed to either mutualistic or pathogenic fungi, regardless of the specific molecular signals involved, we treated rice roots with the complete mixture of native exudates from either the mutualistic *Rhizophagus irregularis* or the pathogenic *Magnaporthe oryzae* fungus, the causal agent of the rice blast disease, and sequenced the roots’ transcriptional response. We compared these transcriptomes to early symbiotic engagement with *R. irregularis* and the well-characterised immunogenic non-fungal epitope flg22 for symbiotic and immunity activation, respectively. While we failed to identify a distinct transcriptional signature associated with exudates of either the beneficial or the pathogenic fungus, we documented a commonly intensified general stress response and PTI related response, accompanied by suppression of critical symbiosis genes. However, when the roots initiated physical engagement with the AM fungus, a dramatic reversal of the root’s transcriptional response was observed, now supporting symbiosis development. Our study therefore reveals that additional checkpoints during pre-symbiosis, possibly at the stage of physical contact, are critical for the recognition of beneficial fungi.

## Results

### Establishing reference transcriptional signatures for early symbiotic interaction and PTI

To set the framework for transcriptome data from rice roots exposed to germinated spore exudates (GSE) from mutualistic or pathogenic fungi, we established reference symbiotic and immunogenic root transcriptomes, using early stages of root interaction with *R. irregularis* (hereafter ‘free AMF’) and flg22 treatment, respectively. At four weeks post-inoculation (wpi), early arbuscular mycorrhizal (AM) symbiosis was established, with ∼20% total root length colonisation; and only ∼5% arbuscule colonisation (figure S1). A total of 164 differentially expressed genes were identified (DEGs; |FC| > 1.5, adjusted p-value< 0.05), including well-described ‘early’ symbiosis marker genes for AM such as *AM1* and *AM3* (31). Markers for the stage of arbuscule development were detectable, including those for symbiotic nutrient exchange, such as *RAM1*, *STR1,* or *PT11* (figure 1A), also reflected by the activation of relevant Gene Ontology (GO) terms including phosphate ion transport and lipid biosynthesis (figure S2A). To frame the stage of symbiotic engagement, we next compared our results with published data from a fully established symbiosis between rice and *R. irregularis* by Das et al., 2022 (32). We found that at our early timepoint of four wpi, only a subset of the AM-induced transcriptome (21 genes) was significantly activated, while 98 genes (e.g., *STR2, NOPE1, HA1*) were strongly induced later at six wpi (figure 1A). Some of these later-stage genes showed mild upregulation at four wpi, but their expression levels were below the threshold to be classified as DEGs (figure S2B). This confirms that our data captured an early stage of symbiosis development, and was therefore considered an appropriate comparative control.

**Figure 1.**
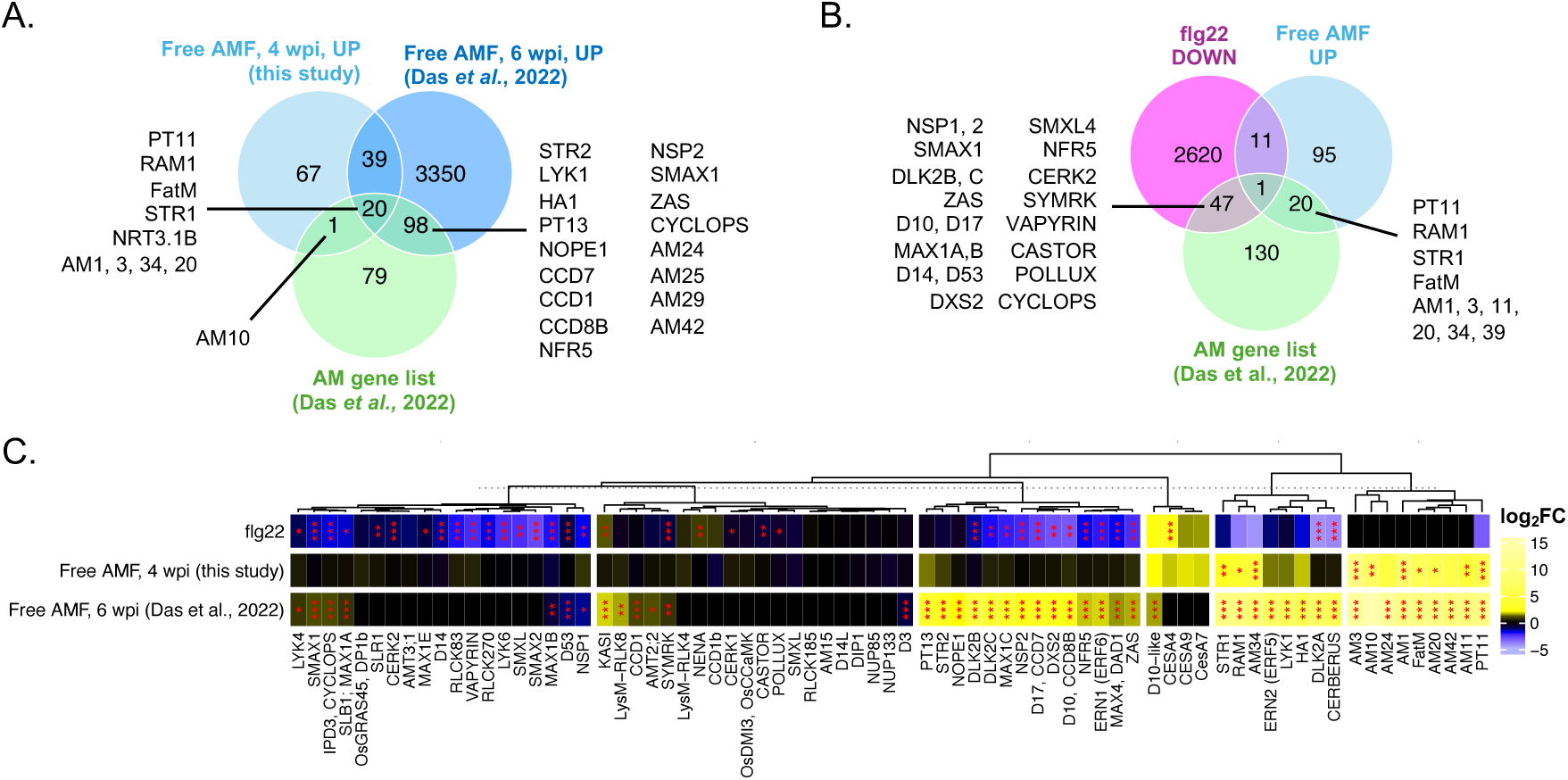
Transcriptional signatures for early symbiotic signalling and PTI activation. (A) Venn diagram of overlaps between up-regulated for free AMF against the mock (water) control in this study after four weeks of co-cultivation, from Das et al., 2022 (32) at a later time-point (six weeks), and the list of AM-related genes in rice from (32). Selected genes of interest highlighted. Venn diagram of overlaps between down-regulated genes for flg22 treatment against the mock (water) control, up-regulated for free AMF, and the list of AM-related genes in rice from (32). Selected genes of interest highlighted. (C) Heatmap of log_2_ fold-changes (log_2_FC) for flg22 and free AMF against water control for the wild-type in this study, as well as the free AMF equivalent treatment from (32) at a later co-cultivation time-point (six weeks), for a selection of genes related to AM symbiosis (common names shown, gene IDs detailed on figure S2B). Significance levels as determined by DEG using DESeq2 shown with asterisks (* for p-value <0.05, ** for < 0.01, *** for< 0.001). Genes were subjected to hierarchical clustering followed by k-means partitioning to subdivide the set in 6 groups, dendrogram show above.

For immune signalling, we chose flg22 as reference treatment as it is an extensively studied PAMP, potently activating PTI in many plant species, including rice (33–35). The use of a bacterial PAMP as a control was deemed useful by allowing to differentiate general PTI signatures from those specific to fungal MAMPs. Furthermore, root transcriptional responses to flg22, and thus its potential influence on AM-related genetic components, had not been previously examined in rice. Treatment with flg22 elicited dramatic transcriptional reprogramming, activating 2,278 and downregulating 2,679 genes. This included genes associated with PTI and defence such as such as *WRKY* transcription factors, chitinases, and other pathogenesis-related (PR) proteins (figure S3A, B). GO enrichment analysis highlighted significant activation of pathways related to hydrogen peroxide, jasmonic acid, salicylic acid and translation (figure S3C), which are hallmarks of leaf immune responses, as is observed in rice during fungal infection by *M. oryzae,* or also in both Arabidopsis and rice in response to MAMPs (33, 35–37). This response was corroborated by comparisons with prior studies of flg22 or pathogen treatments on rice shoot or seedlings (35–38) showing substantial overlaps (figure S3A). Differences in the datasets probably reflect variations in experimental conditions (3h for flg22 by Tang et al., 2021) or plant tissues examined (seedlings for Tang et al., 2021, leaves for Yang et al., 2021). The largest overlap is that with *M. oryzae* infected rice leaves (36), sharing 208 DEGs, including a clear set of defence markers such as *PR1a*, *PR1b*, *RBBI3-1*, *CPS4* and *WRKY45* (figure S3A, B). Together, we conclude that flg22 treatment is a suitable reference for immunity signalling in rice roots. In addition, we unexpectedly found that flg22-triggered PTI resulted in the suppression of a suite of key genes critical for symbiosis signalling (e.g. *NFR5, NSP1, NSP2, VAPYRIN, CYCLOPS, RAM1*) as well as key metabolic enzymes in apocarotenoid hormone biosynthesis and signalling (e.g. *DXS2, D17, D10, MAX1a,b,c, MAX4, ZAS, DLK2B*, *DLK2C)* (figure 1B, C). In rice roots, PTI activation therefore downregulates multiple aspects of symbiosis signalling including receptors, transcriptional regulators, and associated metabolic pathways. This provides a novel insight into the mechanisms by which PTI suppresses symbiosis signalling in a model host species capable of forming AM associations.

### Transcriptional responses of rice roots to fungal exudates from beneficial and pathogenic fungi are highly overlapping

To capture the full complement of known and unknown fungal signals at physiologically relevant concentrations for signalling, we collected whole GSEs from the AM fungal model *Rhizophagus irregularis* and from the fungal rice blast pathogen *Magnaporthe oryzae.* The later infects rice roots with a biotrophic lifestyle, resulting in pathogenic symptoms such as black lesions, as well as induction of immune responses and transcriptional signatures (37, 39–41). Therefore, *M. oryzae* was deemed an appropriate non-symbiotic root-infecting fungus whose exudates would potentially elicit a contrasting transcriptional response to *R. irregularis*. Equivalent to prior studies with *R. irregularis* GSEs on rice (42) and treatments with chitins in rice and other plant species (9, 33, 35, 43–45), roots of four week-old wild-type rice were exposed to GSEs for six hours. Rice roots significantly responded to exudates from water-germinated fungal spores (‘naïve GSE’) of either fungus, corresponding to 495 DEGs for *R. irregularis* and 696 for *M. oryzae*. An unexpectedly large proportion of 313 DEGs were shared between the transcriptomes, corresponding to 63% and 45% of the overall response to GSE from *R. irregularis* and *M. oryzae*, respectively (figure 2A). Only 24 DEGs were shared between *R. irregularis* GSE and the early symbiotic stage (‘free AMF’), of which the majority of two thirds were also shared with *M. oryzae* GSE (figure 2A). No clear activation of known symbiotic genes was observed for *R. irregularis* GSE (figure S2B). Rather an overlap with flg22 treatment was observed for 247 DEGs and 351 DEGs after treatment with *R. irregularis* GSE and *M. oryzae* GSE, respectively, with 171 of these being common to both fungal GSEs (figure 2B). Amongst these, a set of symbiosis and apocarotenoid hormone associated genes (*NFR5*, *NSP2, MAX1C* and *DLK2B* ) was commonly repressed by both fungal GSEs and flg22 treatment (figure S2B). Overall, this suggests broad commonalities in the perception of the different fungal GSEs, in a non-symbiotic manner more similar to PAMP/MAMP responses, as triggered by flg22. In line with this, the most significantly enriched GO terms included response to chemical, toxin catabolic process and glutathione metabolic process, for both the *R. irregularis* and *M. oryzae* GSE treatments (figure S4).

**Figure 2.**
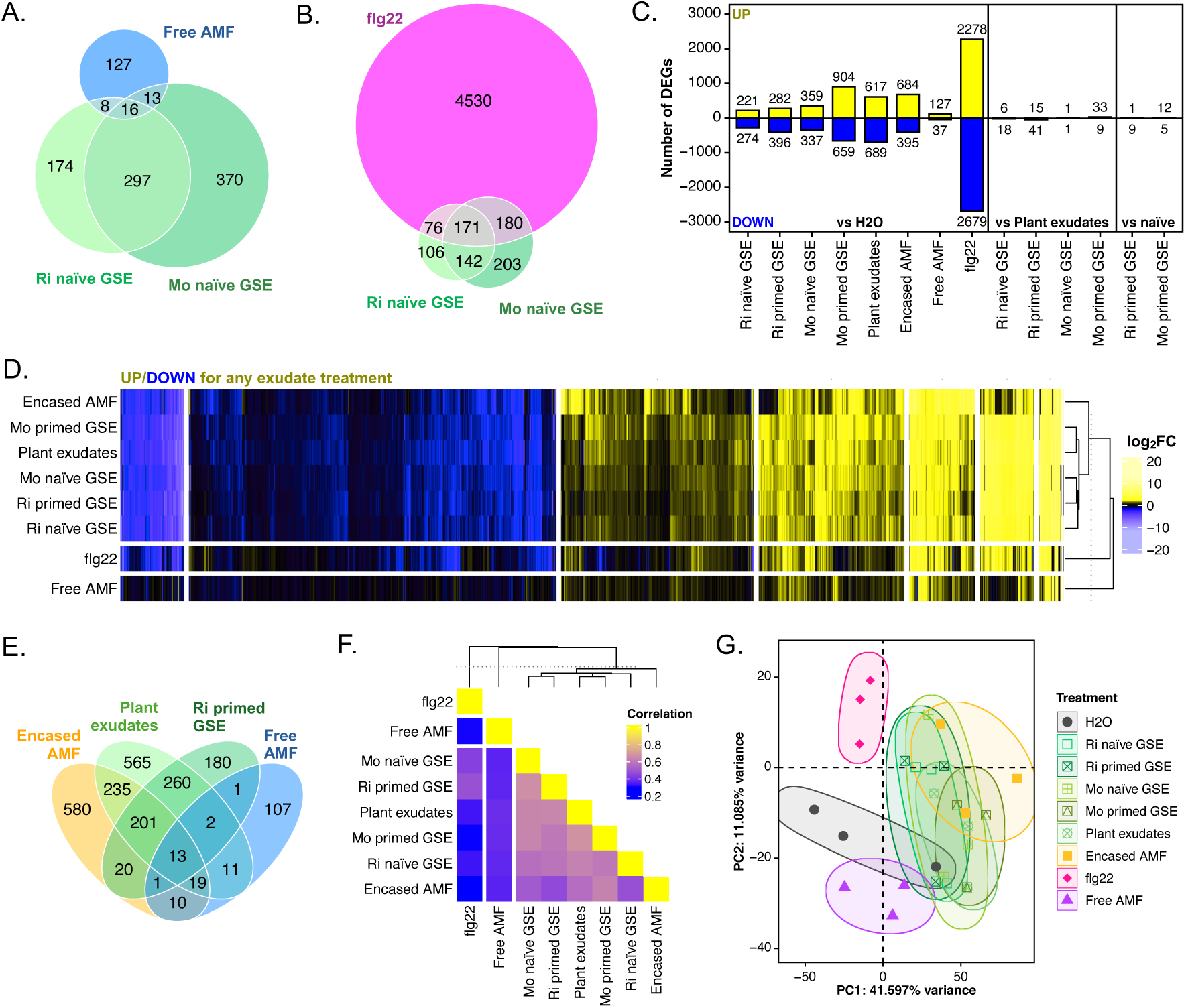
Transcriptional responses of rice roots to exudates are surprisingly similar regardless of their origin, priming or treatment duration. (A, B) Euler diagram of overlaps between DEGs for free AMF (A) or flg22 (B), *R. irregularis* and *M. oryzae* naïve GSE treatments against the mock water control. Area of sections proportional to the number of genes in each set. Number of differentially expressed genes (DEGs), both up-regulated (positive values, yellow) and down-regulated (negative values, blue), for treatments of wild-type rice with free AMF, flg22 and exudate treatments, first all compared against the mock water control, then GSE against plant exudates, then primed GSE against naïve GSE. Numbers over or below bars indicate the precise number of up or down-regulated genes in each comparison. (D) Heatmap of log_2_FC for all DEGs for any exudate treatment against the water control. Both treatments and genes were subjected to hierarchical clustering followed by k-means partitioning to subdivide the set in seven groups for genes and three for treatments, dendrogram for treatments shown on the right, omitted for genes due to high numbers. (E) Venn diagram of overlaps between DEGs for encased AMF, *R. irregularis* primed GSE, plant exudates and free AMF treatments against the mock water control. (F) Heatmap of Pearson correlation coefficients calculated between the log_2_FC for all genes of comparisons of treatments against the water control. Rows and columns were subjected to hierarchical clustering followed by k-means partitioning to subdivide the set in three groups, dendrogram show on top. (G) Principal Components Analysis (PCA) for RNA-seq samples of wild-type rice roots exposed to water (circles, grey), exudates (empty squares, green shades, differing for each), encased AMF (full squares, yellow), free AMF (triangles, violet) or flg22 (rhombuses, pink). Variance-stabilised counts (VST counts) for the 2000 top variable features in the dataset were used to conduct PCA analysis. Only PC1 and PC2 are shown as they together account for more than 50% of the variance. Best fit ellipses for each treatment group were created using the Khachiyan algorithm from the ggforce package in R.

The DEGs were compared to 141 DEGs of a similarly conducted previous report sampled over five time points up to 24h (42), revealing 29 (20.6%) are in common with both *R. irregularis* and *M. oryzae* GSE (figure S5A). The robustness of the response between experiments becomes clearer when looking at fold-changes (most genes up or down-regulated for the earlier study following the same pattern for our dataset, albeit under the threshold to be considered DEGs, figure S5B) or correlation (R^2^>0.3, p-value<0.001, figure S5C), further corroborating the relatedness of the response to exudates from either fungus, reflecting broad commonalities in the perception of the different fungal GSEs.

### Priming beneficial or pathogenic fungi with plant exudates does not alter the plant’s transcriptional response to GSE

We reasoned that plant priming of the fungi in the rhizosphere might be necessary to induce the release of specific and distinct signals from each fungus. Previous research has demonstrated that arbuscular mycorrhizal fungi (AMF) exhibit increased production of short-chain chitin oligomers (CO4-5) when exposed to plant exudates or synthetic strigolactone GR24 (12, 46). We therefore treated germinating fungal spores from *R. irregularis* and *M. oryzae* with root exudates collected from six week-old rice plants grown under phosphate starvation conditions without fungal inoculation (‘primed’ GSEs). The magnitude of transcriptional responses of rice roots treated with primed GSEs were generally higher than to naïve GSEs, with 678 DEGs for *R. irregularis* and 1079 for *M. oryzae*. However, the control treatment – of plants responding to just plant exudates – also led to 1306 DEGs (figure 2C). When comparing each primed GSE treatment to the plant exudate control, fewer than 60 DEGs were identified in both cases. In the primed versus naïve GSE comparisons, only 10 and 17 DEGs were found for *R. irregularis* and *M. oryzae,* respectively (figure 2C). These results indicate highly similar transcriptional responses to either fungus, regardless of priming, and also relative to plant exudates.

Performing k-means clustering of DEG fold-changes between each exudate treatment against water revealed the absence of any gene cluster specific to any individual exudate, plant or fungal. Instead, all exudate treatments grouped together and were clearly distinct from both free AMF and flg22 treatments (figure 2D). In line with this, GO terms enriched in both primed and plant exudate treatments were also considerably overlapping, including “response to chemical” and “glutathione metabolic process” (figure S4). When comparing exudate treatments to the free AMF transcriptome and examining known genes associated to AMS, no activation of symbiotic-specific genes was observed in the *R. irregularis* primed GSE. Instead, there was a significant repression of genes associated with early signalling such as *NFR5*, *NSP2* and enzymes involved in apocarotenoid hormone biosynthesis, a pattern shared across all exudate treatments and related to flg22 treatment (figure S2B). Similarly, we observed shared fold-change trends and enriched GO-terms – such as “response to chemical” or “jasmonic acid” (figure S4) – as well as partial activation of defence markers observed during *M. oryzae* leaf infection (figure S3B). However, these responses were less pronounced than those triggered by flg22. This together suggests that the “common exudate response” involves signalling and transcriptional responses reminiscent of PTI, albeit at a reduced magnitude compared to the flg22-triggered response.

### Prolonged reciprocal signalling between rice roots and AMF reproduces generic transcriptional signature

To establish an experimental set-up that most closely resembles native pre-symbiotic signalling exchanges with free-living AM fungi, *R. irregularis* spores were encased in a nylon membrane physically inaccessible to rice roots for a prolonged period of four weeks of co-cultivation. The transcriptional response to encased AMF resulted in a total 1079 DEGs, similar in magnitude to the exudate treatments. These DEGs overlapped substantially with those induced by GSE and plant exudate treatments, namely 468 DEGs were shared with the plant exudate response, and 235 overlapped with *R. irregularis* primed GSE response (figure 2E). In contrast, only 43 DEGs were shared with the free AMF condition, of which 75% were also differentially expressed in response to plant exudates (figure 2E). Noteworthy, most DEGs for both long and short-term exudate treatments displayed similar expression patterns regardless of treatment, although distinct from the responses to free AMF and flg22 (figure 2D). Examining Pearson pairwise correlations revealed that the transcriptional response to encased AMF was more similar to the exudate treatments (coefficients above 0.45, average 0.53) than to free AMF or flg22 (coefficients of 0.36 and 0.26, respectively, figure 2F). Principal Components Analysis (PCA) further corroborated the lack of discrete transcriptional profiles, as all exudate treatments (including encased AMF) exhibited overlapping clusters. It also revealed a distinct separation from free AMF where plant and fungus started to engage, flg22 treated samples, and the water control (figure 2G). Furthermore, this collective exudate-elicited transcriptional response includes the repression of symbiosis-associated genes and partial activation of defence signatures described earlier (figure 3A, S3B), and shares similar GO term enrichment to exudate treatments, such as response to jasmonic acid, chemical and glutathione metabolic process (figure 3B, S4). The high similarity between encased AMF and GSE treatments supports this non-specific response to exudates being biologically relevant and not artefactual, considering the distinct experimental conditions: 4-week exposure to exudates from spores naturally germinating in a nylon pouch, versus 6h exposure to fungal spore exudates collected in axenic liquid media. Overall, these findings are consistent with the absence of specific pre-symbiotic fungal signals, while suggesting the occurrence of a more distinct transcriptional shift upon physical contact.

**Figure 3.**
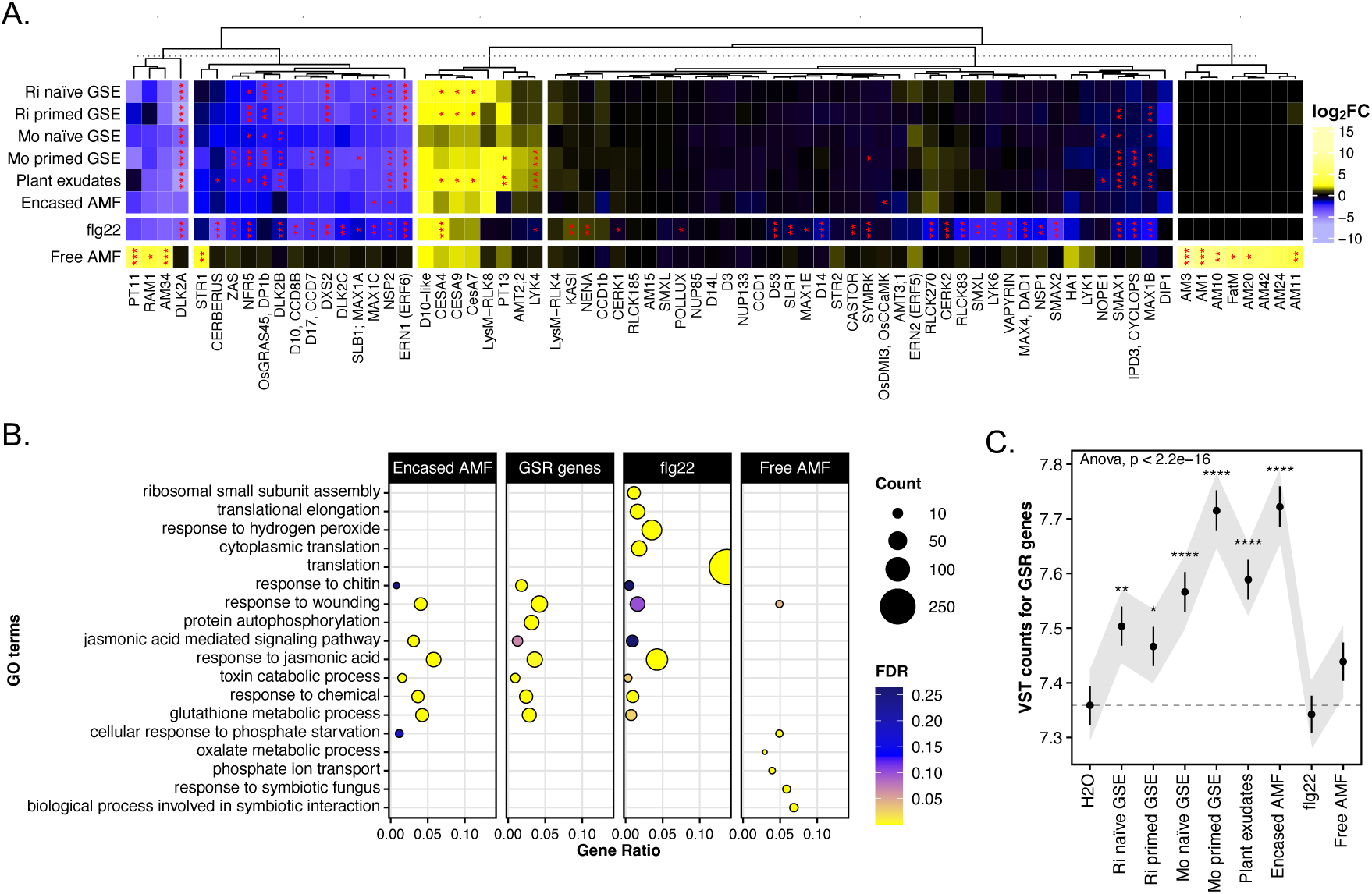
Common transcriptional response of rice roots to exudates features repression of symbiosis-related genes and activation of a general stress response. (A) Heatmap of log_2_FC for exudate treatments, flg22 and free AMF against water control for the wild-type in this study, for a selection of genes related to AM symbiosis (common names shown, gene IDs detailed on figure S2B). Significance levels as determined by DEG using DESeq2 shown with asterisks (* for p-value <0.05, ** for < 0.01, *** for < 0.001). Genes were subjected to hierarchical clustering followed by k-means partitioning to subdivide the set in five groups, dendrogram show above. (B) GO term enrichment for up-regulated genes in encased AMF, free AMF, flg22 against water control, and rice orthologues of General Stress Response (GSR) genes found by Bjornson et al., 2021 (33) in Arabidopsis; top five enriched GO terms by gene ratio for each comparison shown. Colour scale indicates significance as q-value or FDR; dot size proportional to the number of genes in each set included in the GO term; x-axis indicates the ratio of genes up-regulated in each set included in the GO term to the total number of genes annotated with the GO term. (C) Average VST counts plot for relevant wild-type samples for rice orthologues of General Stress Response (GSR) genes, dots indicate average VST counts, errorbars indicate standard error, shadows represent the normal-based 95% confidence interval around the mean. An ANOVA test was performed followed by pairwise t-test between each treatment and the H2O control, significance levels shown with asterisks (* for p-value <0.05, ** for < 0.01, *** for < 0.001 and **** for <0.0001).

Due to the non-specific and PTI-related signature of the exudate response, we hypothesised that this response – whether after short- or long-term exposure – largely comprises the General Stress Response (GSR), a rapid and transient induction of a core set of genes in response to various stimuli, including MAMP and PAMPs in Arabidopsis (33). In support, rice orthologues for a set of ∼1000 Arabidopsis genes known to be induced by MAMP-triggered GSR (33) were significantly upregulated in exudate and encased AMF treatments, but not in free AMF and flg22, and also showed an enrichment for similar GO terms as GSE and encased AMF treatments (figure 3B, C). Together, these findings pinpoint that the shared exudate response represents a general stress-like response, explaining its non-specific nature in contrast to the defined symbiotic and immune transcriptional signatures.

### AM mutants show wild-type response to encased AMF and an impaired response to free AMF

To further investigate if pre-contact fungal signalling was indeed generic and contact formation was critical for specificity, we utilised a panel of plant mutants impaired at various stages of AM interaction. Mutants in this panel were exposed to either prolonged fungal signalling (encased AMF), or inoculated with *R. irregularis* allowing physical contact (free AMF). These mutants span a range of colonisation phenotypes from (a near) complete lack of contact formation as observed in *d14l* (42, 47) to abolished cortex invasion for the CSSP mutants *pollux*, *ccamk* and *cyclops* (31, 42), to quantitatively reduced colonisation for LysM receptor mutants *cerk1, cerk1/2* and *nfr5* (16, 17, 19, 48) (figure 4A, S1).

**Figure 4.**
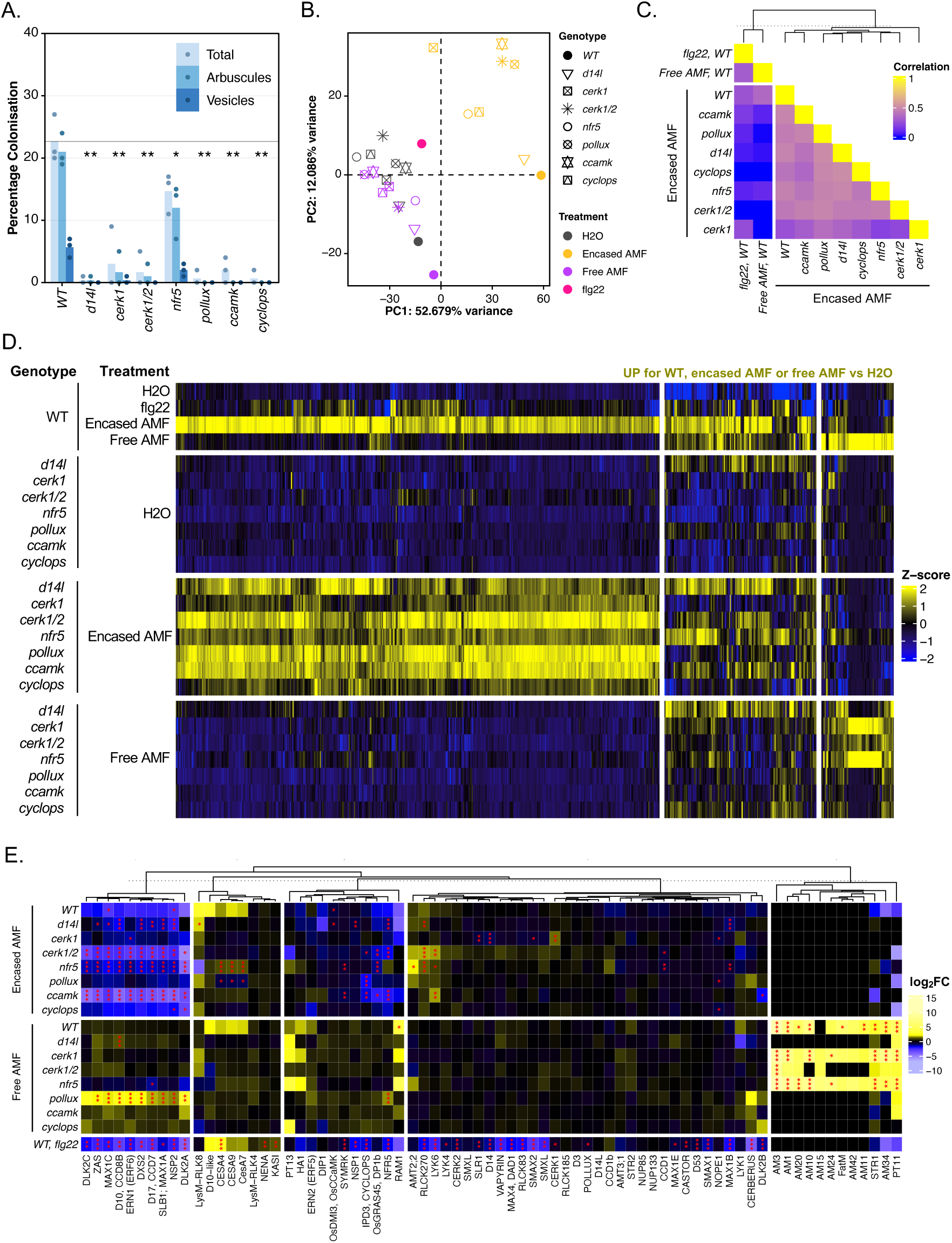
Mutants in AM signalling respond to encased AMF similarly to the wild-type, while their response to free AMF is impaired. (A) Root length colonisation of all rice genotypes at 4 weeks-post-inoculation, each bar indicates the average percentage value of the fungal structure (only total, arbuscules and vesicles shown, the rest detailed in figure S1), all biological replicates are shown, each as individual dots; Kruskal-Wallis test was performed to compare total colonization across genotypes, resulting in a 0.03813 p-value, under the 5% significance level, followed by one-sided t-test between each treatment and the wild-type (WT) control, significance levels shown with asterisks (* for p-value <0.05, ** for < 0.01, *** for < 0.001 and **** for <0.0001). (B) Principal Components Analysis (PCA) for RNA-seq samples of wild-type and AM signalling mutants (genotype indicated by point shape) roots when exposed to H2O (grey), encased AMF (yellow), free AMF (violet) or flg22 (pink). Variance-stabilised counts (VST counts) for the 2000 top variable features in the dataset were used to conduct PCA analysis, only PC1 and PC2 are shown as they together account for more than 50% of the variance. VST counts of replicates for the same genotype and treatment were averaged to facilitate visualization. (C) Heatmap of Pearson correlation coefficients calculated between the log_2_FC for all genes of comparisons of treatments shown against the water control for each genotype. Rows and columns were subjected to hierarchical clustering followed by k-means partitioning to subdivide the set in two groups, dendrogram show on top. (D) Heatmap of VST row-centered counts for up-regulated DEGs in encased AMF and free AMF against the H2O control for the wild-type, of all mutants and wild-type exposed to H2O, encased AMF or free AMF, also flg22 for the wild-type. Genes were subjected to hierarchical clustering followed by k-means partitioning to subdivide the set in 3 groups, dendrogram omitted. (E) Heatmap of log_2_FC for free and encased AMF treatments against water for wild-type and AM mutants, for a selection of genes related to AMS. Significance levels as determined by DEG by DESeq2 shown with asterisks (* for p-value <0.05, ** for < 0.01, *** for < 0.001). Genes were subjected to hierarchical clustering followed by k-means partitioning to subdivide the set in five groups, dendrogram show on the left.

When exposed to physically separated encased AMF for four weeks, all mutants showed a clear and strong transcriptional response with around 1,000 to 3,000 DEGs (figure S6). PCA (figure 4B) and pairwise correlation coefficients (figure 4C) analyses demonstrated that the transcriptional responses of all mutants to encased AMF closely aligned with the wild-type response. Conversely, their transcriptional response to free AMF mirrors the extent of root colonisation: *nfr5*, *cerk1* and *cerk1/2* partially recapitulated the upregulation of the same set of symbiotic genes as the wild-type, a response mostly abolished in the rest of the mutants [*pollux, ccamk, cyclops, d14l;* (figure 4D, E)]. The PCA also showed that the mutants’ response to free AMF was substantially altered: samples clustered more closely together with the wild-type water control or flg22 treatment than with wild-type permitted to engage with free AMF, in part reflecting the reduced transcriptional response and limited AM colonisation levels (figure 4B). This difference is clearer when examining normalised gene counts for activated genes, as most genes upregulated in the encased AMF scenario in the wild-type were also induced in all mutants (figure 4D). Whereas the transcriptional response to free AMF was largely abolished except in mutants that still allowed intraradical colonisation by *R. irregularis* (figure 4D).

Focusing on the suite of symbiosis, defence-related and GSR genes, all signalling mutants showed the same pattern as wild-type in response to encased AMF: repressing symbiosis signalling genes such as *NFR5*, *NSP2*, and butenolide biosynthetic genes (figure 4E), activating PTI-related markers induced by flg22 and *M. oryzae* infection (figure S7) as well as GSR (figure S8A, B). By contrast, when physical contact is permitted in the free AMF condition, suppression of symbiosis signalling genes is relieved in all mutants (figure 4E), accompanied by the simultaneous suppression of immunity genes (figure S7). GSR induced during pre-contact signalling by encased AMF treatment was similarly relaxed in the mutants, regardless of the presence or absence of symbiosis establishment (figure S8A, C). This phenomenon is striking, and demonstrates defined properties for the suppression of symbiosis signalling genes, and the activation of immunity and GSR: (1) it is generic and occurs in early biotic signalling, (2) it is independent of the CSSP and (3) is reversed later when physical contact is permitted, again transcending the common symbiosis signalling pathway.

## Discussion

We initially sought to define transcriptional signatures underpinning symbiotic versus immunity activation during the early, pre-symbiotic phase – when fungal-derived molecules are perceived by the plant. This motivation was shaped by the prevailing, but increasingly nuanced, notion that perception of particular chitin derivatives provide specificity: activating symbiosis with CO4, LCO, CO8 and triggering immunity with long-chain chito-oligosaccharides, e.g. CO8. Meanwhile, there is increasing evidence that other signalling molecules from AM fungi are involved in pre-symbiotic signalling – several of which proteinaceous – to facilitate plant recognition and symbiosis. We therefore reasoned that using the exudates released by the model AM fungi *R. irregularis* will capture the full cocktail of signalling molecules, both known and unknown. Moreover, transcriptomic profiles of plant responses to fungal exudates are relatively limited. In parallel to exudates from symbiotic AM fungi, the pathogenic *M. oryzae* and flg22-activated PTI provided important benchmarks for transcriptional specificity.

In this study, we unexpectedly found a remarkable lack of specificity in the transcriptomic response of rice roots to exudates of a beneficial and a pathogenic fungus. The bouquet of signals released by either fungus failed to induce a specific immunity or symbiotic transcriptional signature. Neither priming the fungi with plant exudates, nor long-term co-cultivation of AM fungi in close proximity *sans* physical contact, produced a differentiated transcriptional response. Instead, common to the set of short- and long-term exudate perception is the suppression of a branch of symbiosis signalling genes (*NFR5*, *NSP2*, butenolide biosynthesis), activation of PTI-related defense markers and a general stress-response signature. A portion of this signature such as suppression of symbiosis signalling genes, in fact, is shared with flg22-activated PTI. Furthermore, this signature was observed in mutants impaired at various stages of AM symbiosis, indicating that it is unlikely to be a symbiotic response, and is not controlled by the canonical CSSP. The third aspect of the transcriptional signature is the substantial upregulation of the non-specific GSR (33, 49), which is not highly induced during AM symbiosis.

In contrast, when AM fungi hyphae are not physically encased but allowed to engage freely with plant roots, specific activation of symbiosis signalling genes occur, alongside the reversal of the exudate response signatures: relief of suppression of symbiosis signalling, partial suppression of PTI/defence-related markers and attenuation of the GSR. While the activation of symbiosis genes required a functional CSSP, the latter two aspects transcended it. How this occurs is currently unknown and lays the foundation for future investigation. The range of genetic mutants utilised here revealed that additional layers of signal exchanges running parallel to the CSSP must exist, and it culminates in a phenotype of stress- and immune-modulation.

Our observations extend earlier work in the understanding of plant colonisation by AM fungi. These previous studies – based on targeted analysis of specific proteins or genes, or broader surveys using microarrays – revealed that early perception of AM fungi often involve activation of stress- and defence-related responses in the roots. These responses are typically attenuated when symbiosis is established (50–54) reviewed in (55). Although, comprehensive gene expression profiling by exposing hosts to fungal exudates of different origins is scarce, existing work also revealed that both pathogenic and AM fungi exudates activate overlapping, and defence-related responses (56). In addition, transcriptomic and cytological analyses in host and non-host plants suggest further signalling steps and processes to be involved in determining compatible (here symbiotic) and incompatible interactions (57–59).

Taken together, our results favour the scenario where symbiosis is not defined during the molecular dialogue in the rhizosphere, or specified solely at the level of chitin perception, but rather require multiple aspects of signal exchanges on top of activating the CSSP. Modulating stress and immune-related responses triggered by early pre-symbiotic signal exchanges is one, and it may be intertwined with the close physical contact between fungal hyphae and plant roots. This scenario opens the possibility for nuanced, sequential signalling steps, for more MAMPs, effectors, DAMPs (damage-associated molecular patterns) to shift the balance between symbiotic and immunity responses before a committed interaction.

## Materials and Methods

### Plant-AM fungal material, plant growth conditions and AM fungi inoculation

All rice experiments were conducted using *Oryza sativa* subsp. japonica cv. Nipponbare (NB) as background genotype. The following mutants were employed: *d14l-3* (T insertion generated by CRISPR-Cas9 (47)), *cerk1-1* (48), *cerk1/2-1* (17), *nfr5-1* [(16); all three generated via homologous recombination], *pollux-3* (31, 60), *ccamk-1* (31, 60) and *cyclops-2* [(31, 60); all three generated via Tos17 insertional mutagenesis (61)]. *O. sativa* seeds were dehusked, surface-sterilised for 1 min in 70% (v/v) ethanol, then 20 min in 5% (v/v) sodium hypochlorite, rinsed with reserve-osmosis water (RO-H_2_O) and germinated in petri dishes containing 0.6% (w/v) Bacto^TM^ Agar sealed with Micropore^TM^ tape in a 30°C incubator in the dark for 7 days. Seedlings were then transferred into cylindrical pots (11 cm depth and 8 cm diameter, 3 seedlings in each) containing autoclaved quartz sand in walk-in phytochambers with photoperiod of 12-hour day-night cycle at 28/20°C, 65% relative humidity under LED illumination at 300 μmol/μm^2^s. The AM fungal model species *Rhizophagus irregularis* (DAOM197198) was employed for all inoculation assays. Free AMF pots were inoculated with 1000 spores of *R. irregularis* per pot, while for encased AMF pots a single closed 0.45 μm nylon membrane (Whatman) filled with sand inoculated with 3000 spores was placed half-covered in sand in the middle of the pot (between the three plantlets). The source of AM fungi innocula was a mix of *Agrobacterium rhizogenes* transformed carrot hairy root cultures (*Daucus carota* L.)(62) from which spores were extracted as previously described (31), and commercial axenic DAOM197198 inoculum (Mycorise® ASP, Premier Tech Biotechnologies, Canada) in a 2:1 ratio. Plants were watered thrice weekly, the first week post inoculation (wpi) with RO-H_2_O, followed by low Pi fertilization twice a week with half-strength Hoagland’s solution containing 25 μM KH_2_PO_4_. These growth conditions were demonstrated previously to promote efficient and equal mycorrhization (20, 31). For the encased AMF pots, the resulting filtered liquid was reapplied to the sand to expose the enclosed fungi to plant exudates (repeated twice each watering).

### Generation of plant exudates and Germinated Spore Exudates (GSE) from R. irregularis and M. oryzae

Plant exudates were collected by growing *O. sativa* plants without AM inoculation in the same conditions as previously described for 6 weeks, then taking the plants out of the pots, rinsing sand off with RO-H_2_O, and incubating 3 plants in 10 mL of half-strength Hoagland’s solution containing 25 μM KH_2_PO_4_ in 60 mL plastic containers (VWR International Ltd) O/N for 16h at 100 rpm in a shaker in the same growth chamber. The resulting solution was filter sterilised, frozen in liquid nitrogen and stored at -80°C. *R. irregularis* DAOM197198 spores were sourced from carrot hairy root cultures as previously described and commercial axenic DAOM197198 inoculum from PremierTech in a 2:1 ratio. *M. oryzae* spores of Guy11 strain expressing eGFP under the *ToxA* promoter (63) were sourced from subcultures. This strain was a generous gift from Nick Talbot (The Sainsbury Laboratory, Norwich). Spores were propagated by growing the subcultures for 2 weeks at 200 µmoles/m^2^/s light, 60% humidity, 16h light/8h light cycle, 28°C in light and 24°C in dark on complete medium as described in Talbot et al., 1993. The medium was covered with small square pieces of Whatman filter paper, which were collected as *M. oryzae* stocks for future subculture, and stored at -20°C, a temperature at which the fungus is metabolically inactive. Spores for GSE production were collected by washing the surface of the subculture medium after 2 weeks of growth with sterile RO-H_2_O, then filtered through a miracloth (GLife Tech, Singapore). After a centrifuge step, *M. oryzae* spores in the pellet were resuspended in sterile RO-H_2_O and the concentration was determined by counting spores in a hemocytometer. To prepare GSE, spores were placed in 1L flasks containing *R. irregularis* liquid growth media (62) supplied with either 2x diluted plant exudates (for primed GSE) or half-strength Hoagland’s containing 25 μM KH_2_PO_4_ (naïve GSE), diluting spores to a final concentration of 1000 spores/mL for *R. irregularis* or 10 000 spores/mL for *M. oryzae*. For *R. irregularis* GSE, the flasks were placed in an incubator at 30°C, dark, 5% CO2 (injected into flask at 0.5 lpm), and 100 rpm, for 4 days. For *M. oryzae* GSE, the flasks were placed in a growth cabinet (licensed for *Magnaporthe* work) with a fluctuating temperature of 28/24°C, 60% relative humidity, in the dark for 4 days. The resulting GSE were filter sterilised using a 0.22 μm filter (Millipore Stericup® Quick Release), frozen in liquid nitrogen and stored at -80°C.

### Treatments with flg22 or GSE before harvesting

Plants were taken out of the pots, sand rinsed off the roots with RO-H_2_O, then left overnight (*c.*16h) in 10 mL of half-strength Hoagland’s solution containing 25 μM KH_2_PO_4_ in 60 mL plastic containers (VWR International Ltd) in the same growth chamber. After this incubation, the Hoagland’s solution was removed, then 10 mL of the appropriate treatment was added: flg22 (10^-6^M, Genscript, USA), GSE, or RO-H_2_O. This results in GSE from an equivalent of 10,000 *R. irregularis* or 100,000 *M. oryzae* spores added to each pot (3 plants). As the appropriate control for GSE, plant exudates were diluted 2x in *R. irregularis* growth media before treatment. After 6 hours of treatment, roots were rinsed, dried, cut and the tissue flash frozen in liquid nitrogen. To minimize impact of circadian clock, all harvests were completed within a span of 2 hours (1200-1400). Tissue was stored at -80°C before RNA extraction.

### Root Staining and Mycorrhizal Quantification

Roots were harvested at 4 wpi. Trypan blue (Sigma-Aldrich) staining and mycorrhizal colonization of the different genotypes were quantified as described previously (20, 31). Quantification of fungal colonization took place by counting extraradical hyphae, intraradical hyphae, hyphopodia, arbuscules, vesicles and spores along 100 different visual fields under a 20× magnification objective using using DM750 Microscope (Leica, Germany) and expressed as percentage of the total root length scored.

### RNA Extraction, cDNA Synthesis, and Gene Expression Analysis

Roots harvested were frozen in liquid nitrogen and stored at -80°C until used for RNA extraction. Root tissues were homogenized with metal beads using Geno/Grinder® (SPEX SamplePrep, USA) at 1500 strokes/minute for 1 min until fine powder. RNA was extracted from ground tissue with TRIzol reagent (Invitrogen, USA) with chloroform washes and isopropanol precipitation, and residual DNA removed by treatment with DNaseI (Invitrogen, USA) following manufacturer’s instructions. RNA integrity and purity were assessed by presence of clear 28S and 18S rRNA bands following gel electrophoresis, and quantity determined by NanoDrop 2000 Spectrophotometer (Thermo-Fisher Scientific, USA). cDNA synthesis was conducted as described previously (20) where reverse transcription was performed on 1.1 μg of RNA using Superscript IV reverse transcriptase and oligo(dT)15 primers following manufacturer’s instructions. Absence of contaminating genomic DNA was confirmed by performing PCR with primers on two exons flanking a spliced intron in GAPDH to yield a smaller product following electrophoresis on a 0.8% (w/v) agarose gel, with gDNA sample as positive control. Quantitative polymerase chain reaction (qPCR) was performed as described previously (20, 31), using SYBR Green Fluorophore on C1000 Thermal Cycler with CFX384 real-time detection system (Bio-Rad, USA). Specific primers (table S1) were used for *CYCLOPHILIN2*, *ACTIN* and *UBIQUITIN* as housekeeping genes, for *AM1*, *AM3*, and *PT11* as rice AM marker genes to validate RNA-seq results (figure S9). Gene expression values were normalized to the geometric mean of the three reference genes and displayed as a function of CYCLOPHILIN2 mRNA levels.

### RNA-Seq library preparation, sequencing, and data analyses

RNA extraction and quality controls were conducted as previously described. RNA samples were diluted to an equal concentration and sent for Illumina NovaSeq 6000 sequencing to Novogene (Cambridge, UK) with a sequencing depth of 3 Gb (10 million reads). Raw reads were pseudo-aligned to the *O. sativa* Nipponbare reference transcriptome (Os-Nipponbare-Reference-IRGSP-1.0.52)(65) using Kallisto v0.46.1 (66) with default parameters, to obtain count estimates. An average 91.6% of the reads were aligned to the rice transcriptome. As a quality control we used STAR v2.7.10b (67) to map the entire genome (including organelles), with default parameters, for a subset of samples. This approach resulted in a very similar alignment percentage (91.87%). Raw reads were also pseudo-aligned to the *R. irregularis* DAOM197198 transcriptome (from (68), NCBI bioproject PRJNA885267) using Kallisto, resulting in an average mapping rate of 0.19%, 0.28% for the mycorrhizal samples. The count estimates were normalised, filtered (keeping only genes with more than 10 counts among all samples), and subjected to pairwise differential expression analysis between conditions using the R package DESeq2 v1.40.2 (69), with a threshold of fold-change ≥ 1.5 or ≤ -1.5 and Benjamini-Hochberg false discovery rate corrected *P*-value < 0.05. Gene Ontology (GO) and Kyoto Encyclopedia of Genes and Genomes (KEGG) enrichment analyses were conducted using the R package clusterProfiler v4.8.3 (70). The annotations of the genes including associated GO and KEGG terms were collected from various sources: The Rice Annotation Project (RAP)(65) and EnsemblPlants (71) annotations for the Os-Nipponbare-Reference-IRGSP-1.0.52 reference genome, Oryzabase (72), the AM gene list from Das et al., 2022 and KEGG (73–75). Rice orthologues of Arabidopsis General Stress Response (GSR) genes from (33) were identified with Orthofinder v2.5.4 (76), using a set of 20 proteomes from Ensembl Plants (*Amborella trichopoda* AMRT1.0, *Arabidopsis thaliana* TAIR10, *Beta vulgaris* RefBeet 1.2.2., *Brachipodium distachyon* v3.0, *Brassica rapa* 1.0, *Cucumis sativus* ASM407v2, *Glycine max* v2.1, *Hordeum vulgare* Morex v3, *Marchantia polymorpha* v1, *Medicago truncatula* MtrunA17r5.0, *Oryza sativa* IRGSP-1.0, *Physcomitrium patens* V3, *Pisum sativum* v1a, *Populus trichocarpa* v3, *Selaginella moellendorffii* v1.0, *Solanum lycopersicum* SL3.0, *Sorghum bicolor* NCBIv3, *Triticum aestivum* IWGSC, *Vitis vinifera* PN40024.v4, *Zea mays* B73-REFERENCE-NAM-5.0), then using the 1-to-1 orthologue list between *Oryza sativa* and *Arabidopsis thaliana*, all rice genes in the same orthogroup as any Arabidopsis relevant gene were included for the analysis.

### Statistical Analyses

Statistical significance of the differences between genotypes, treatments or conditions was assessed by the parametric ANOVA test or the non-parametric Kruskal-Wallis (if the assumptions of ANOVA were not met by the dataset), at 5% significance level. Student’s T-test or Pairwise Wilcoxon rank sum were then respectively used to identify statistically significant differences between genotypes, denoted in graphs with asterisks (* for p-value <0.05, ** for < 0.01, *** for < 0.001 and **** for <0.0001). Validity of these tests was confirmed by a one-way analysis of variance (ANOVA) test for the log10 transformed data (to ensure equal variance). For all tests, the null hypothesis was rejected with a p-value threshold of 0.05. Every graph displays all data points. The exact statistical test used for each dataset is indicated in the corresponding figure legend. All statistical analyses were conducted using R (https://cran.r-project.org).

## Acknowledgments

We thank the Giles Oldroyd, Jongho Sun (University of Cambridge), and Feng Feng (Oklahoma State University) for sharing elicitors used in this study; Katrin Geisler (University of Cambridge) for access to the CO_2_ incubator; Gizem Kiyak for providing *Magnaporthe oryzae* spores. We are grateful to William Summers and Jeongmin Choi for their contributions to early experimental work that helped shape this project, and to various colleagues at the Crop Science Centre, University of Cambridge for assistance with harvesting and constructive feedback. Research was supported by St John’s College (University of Cambridge) through the Benefactor’s Scholarship to G.F.G. and Research Reimbursement Schemes to G.F.G. and U.P., by the Cambridge Philosophical Society through the Research Studentship Scheme to G.F.G., by the National University of Singapore through the Overseas Postdoctoral Fellowship (NUS-OPF), and Development Grant (NUS-DG) 2021/2022 to C.H.C. Research in U.P.’s laboratory was supported by the Biotechnology and Biological Sciences Research Council Grant BB/V006029/1 and BB/V002295/1.

## Data availability

Raw sequences and gene count matrices generated in this study are available in the NCBI GEO database under the accession GSE28377. R code employed to analyse the data and generate graphics is available in the following github repository:

https://github.com/gabriel-ferreras/Presymbiotic_transcriptional_landscape.

Other data supporting the findings of this article are available in the supplemental information.

