## Supplemental Information for "Defining the pre-symbiotic transcriptional landscape of rice roots"

**Supporting Information for**
### Defining the pre-symbiotic transcriptional landscape of rice roots

Gabriel Ferreras Garrucho, Uta Paszkowski*§, Chai Hao Chiu*§

Crop Science Centre, Department of Plant Sciences, University of Cambridge, 93 Lawrence Weaver Rd, Cambridge, United Kingdom, CB3 0LE.

*Corresponding authors: Uta Paszkowski, Chai Hao Chiu
§shared last author

**This PDF file includes:**

Figures S1 to S9
Tables S1
SI References

#### Supplementary figures and tables

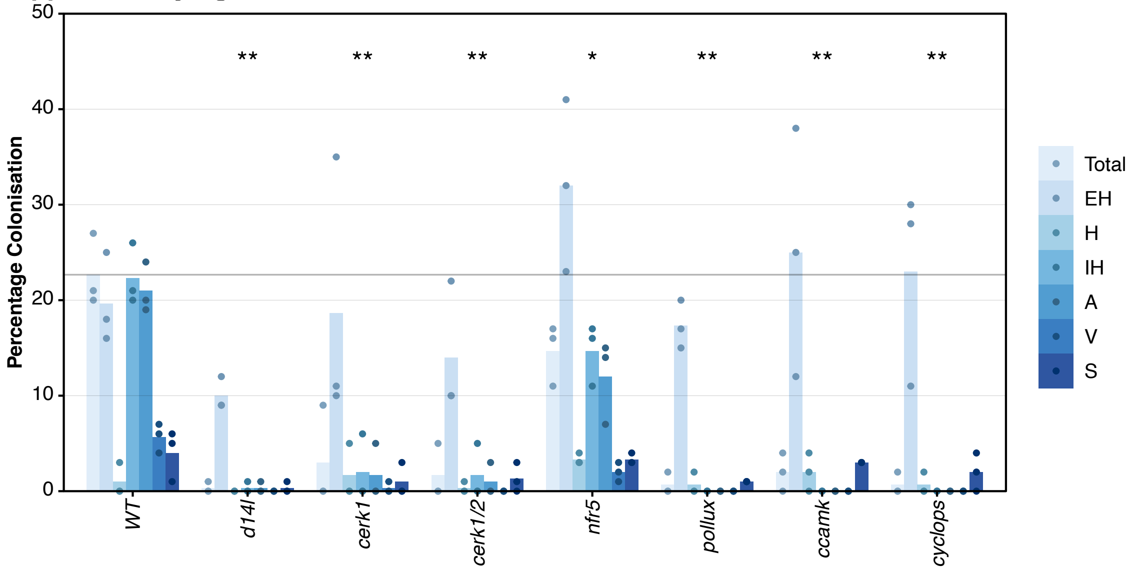


**Fig. S1. Arbuscular mycorrhizal colonisation levels at 4 weeks post-inoculation for all genotypes used in this study.** Root length colonisation determined for all rice genotypes at four weeks-post-inoculation (wpi), each bar indicates the average percentage value of the fungal structure, all biological replicates are shown, each as an individual dot. Kruskal-Wallis test was performed to compare total colonisation across genotypes, resulting in a 0.03813 p-value, under the 5% significance level, followed by one-sided t-test between each treatment and the wild-type control, significance levels shown with asterisks (* for p-value <0.05, ** for < 0.01, *** for < 0.001 and **** for <0.0001). Total, total intraradical colonisation; EH, extraradical hyphae; IH, intraradical hyphae; H, hyphopodia; A, arbuscules; V, vesicles; S, spores.

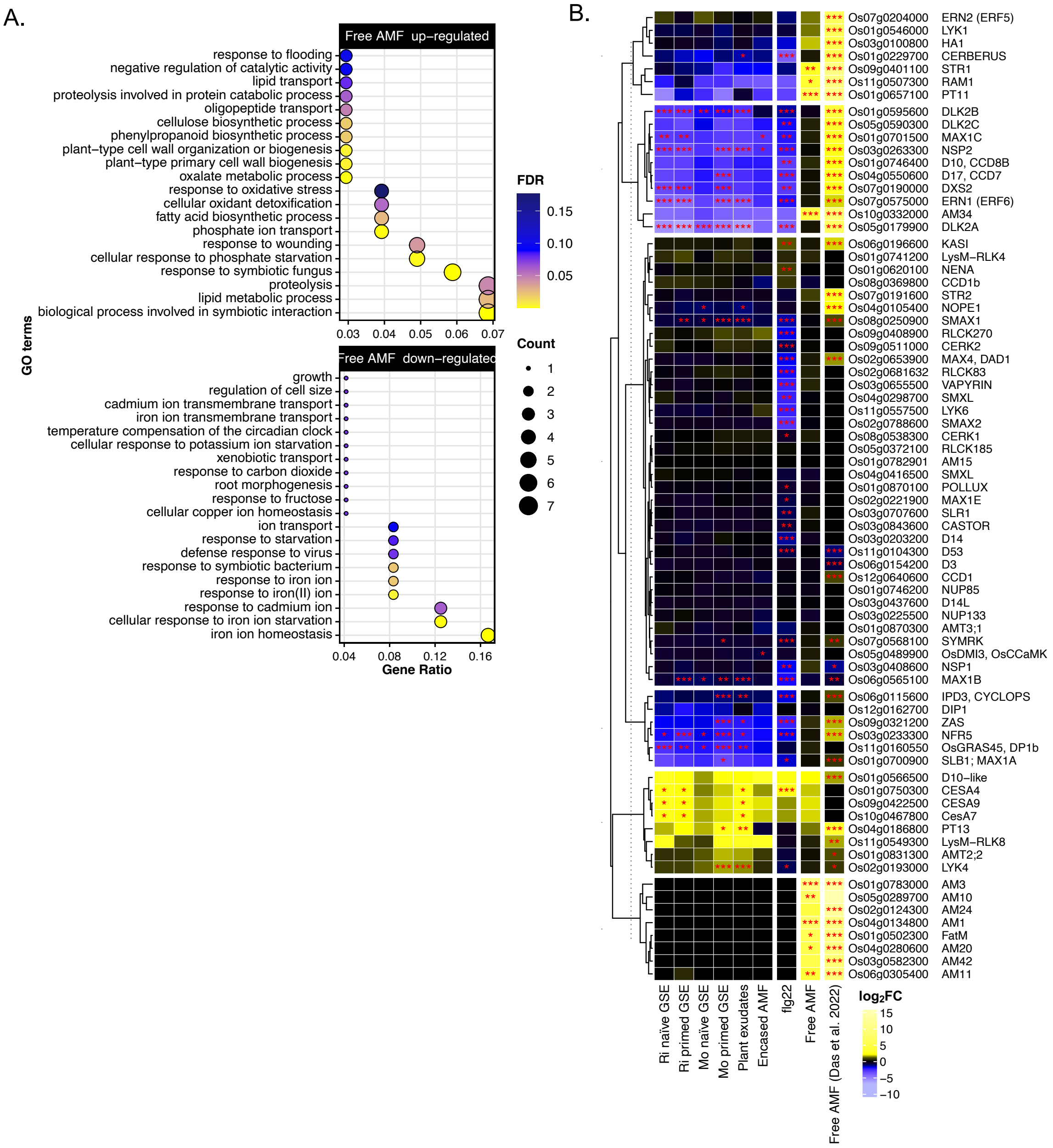


**Fig. S2. Co-cultivation with free AMF for 4 weeks results in a symbiotic transcriptional signature.** (A) GO term enrichment for up and down regulated genes for free AMF against H2O control; top 20 enriched GO terms by gene ratio shown; colour scale indicates significance as q-value or FDR; dot size proportional to the number of genes in each set included in the GO term; x-axis indicates the ratio of genes up-regulated in each set included in the GO term to the total number of genes annotated with the GO term. (B) Heatmap of $\log_2$ fold-changes ($\log_2$FC) for exudate and encased AMF treatments, flg22 and free AMF against H2O control for the wild-type in this study, as well as the free AMF equivalent treatment from Das et al., 2022 (1) at a later co-cultivation time-point (six wpi), for a selection of genes related to AMS. Significance levels as determined by differential gene expression analysis by DESeq2 shown with asterisks (* for p-value <0.05, ** for < 0.01, *** for < 0.001). Genes were subjected to hierarchical clustering followed by k-means partitioning to subdivide the set in six groups, dendrogram show on the left.

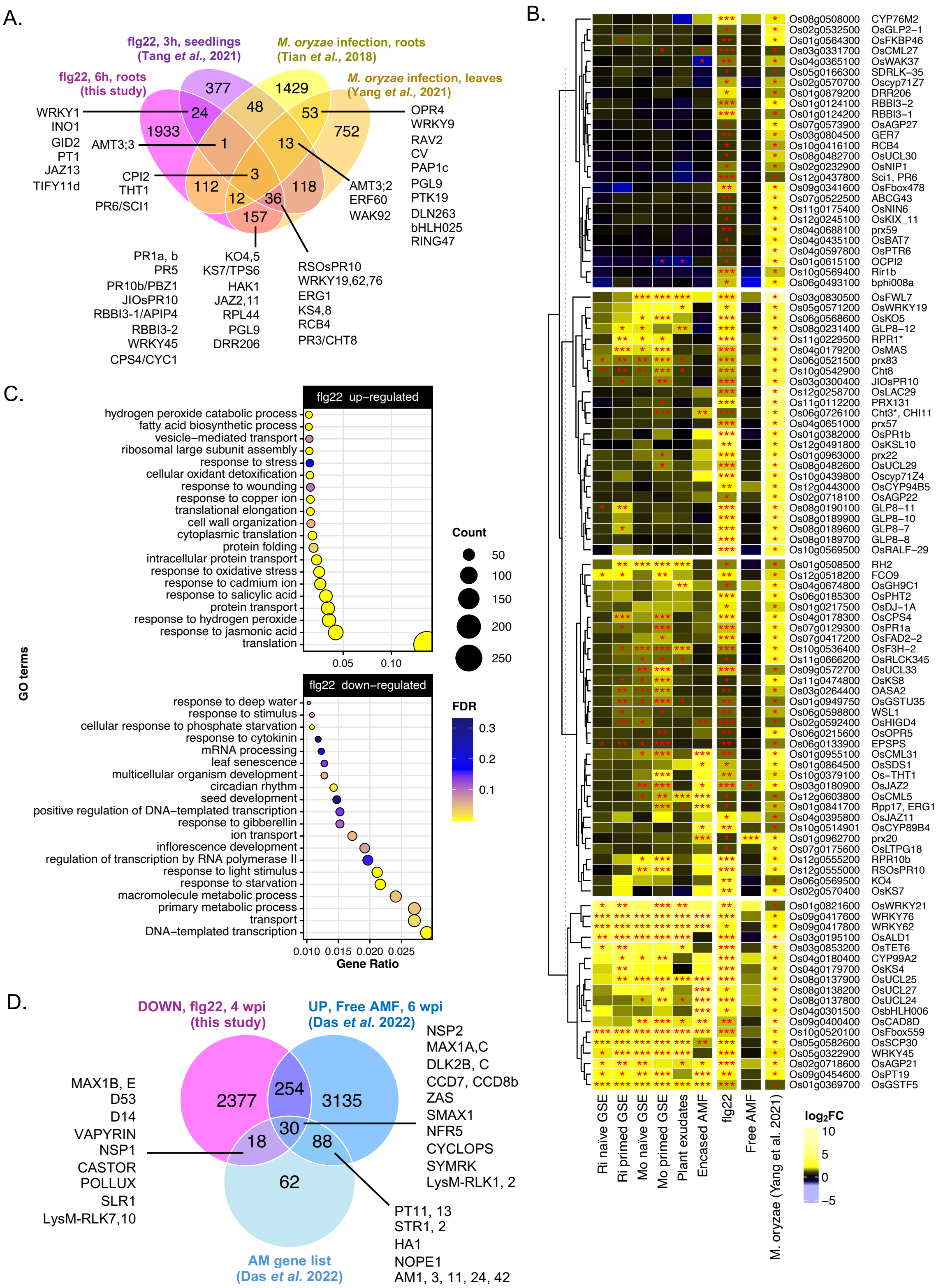


**Fig. S3. Flg22 treatment results in a PTI transcriptional signature.** (A) Venn diagram for overlaps between upregulated genes for flg22 in this study and selected treatments using PAMPs or pathogen infection by *M. oryzae* from the published datasets: seedlings treated with flg22 for 3h (Tang et al., 2021 (2)), *M. oryzae* infected leaves (Yang et al., 2021 (3)), *M. oryzae* infected roots (Tian et al., 2018 (4)). Selected genes of interest highlighted. (B) Heatmap of $log_2FC$ for exudate and encased AMF treatments, flg22 and free AMF against H2O control for the wild-type in this study, as well as *M. oryzae* leaves from Yang et al., 2021, for a selection of genes upregulated both under flg22 treatment in this study and *M. oryzae* infection in leaves (Yang et al., 2021 (3)). Significance levels as determined by differential gene expression analysis by DESeq2 shown with asterisks (* for p-value <0.05, ** for < 0.01, *** for < 0.001). Genes were subjected to hierarchical clustering followed by k-means partitioning to subdivide the set in four groups, dendrogram show on the left. (C) GO term enrichment for up and down regulated genes for flg22 against water control; top 20 enriched GO terms by gene ratio shown; colour scale indicates significance as q-value or FDR; dot size proportional to the number of genes in each set included in the GO term; x-axis indicates the ratio of genes up-regulated in each set included in the GO term to the total number of genes annotated with the GO term. (D) Venn diagram for overlaps between downregulated genes for flg22 in this study, up-regulated under free AMF equivalent treatment from Das et al., 2022 (1) at a later co-cultivation time-point (six wpi), and the list of AM-related genes in rice from Das et al., 2022 (1). Selected genes of interest highlighted.

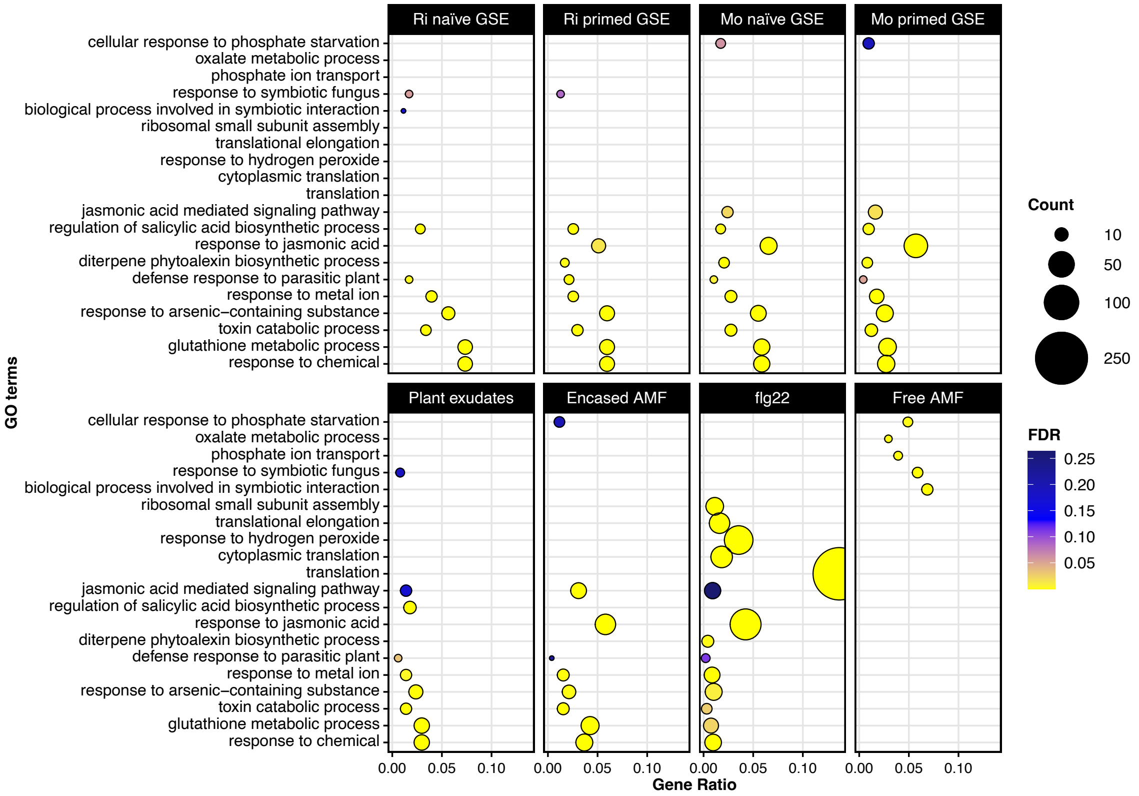


**Fig. S4. Enriched GO terms for up-regulated genes of all exudates treatments are highly overlapping, distinct from free AMF and flg22.** GO term enrichment for up all treatments against water control in the wild-type; top five enriched GO terms by gene ratio for each comparison shown; colour scale indicates significance as q-value or FDR; dot size proportional to the number of genes in each set included in the GO term; x-axis indicates the ratio of genes up-regulated in each set included in the GO term to the total number of genes annotated with the GO term.

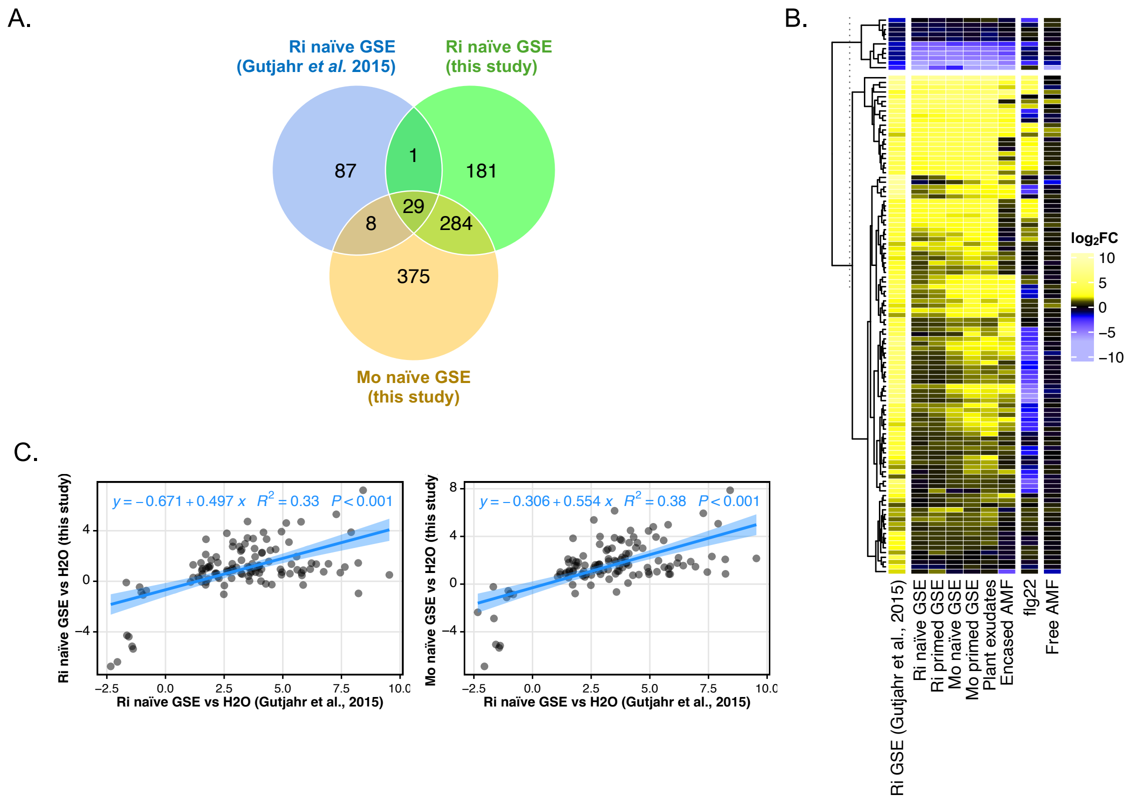


**Fig. S5. Naïve GSE response is replicable between studies but highly overlapping between *R. irregularis* and *M. oryzae* GSE.** (A) Venn diagram for overlaps between upregulated genes for *R. irregularis* naïve GSE treated roots, either in this study or in Gutjahr et al., 2015, as well as *M. oryzae* naïve GSE treated roots. (B) Heatmap of $log_2FC$ for exudate and encased AMF treatments, flg22 and free AMF against H2O control for the wild-type in this study, as well as *R. irregularis* naïve GSE treated roots from Gutjahr et al., 2015 (5), for DEGs from this last dataset. Genes were subjected to hierarchical clustering followed by k-means partitioning to subdivide the set in two groups, dendrogram show on the left. (C) Scatterplots of $log_2FC$ for DEGs in the Gutjahr et al., 2015 (5) *R. irregularis* naïve GSE treatment against naïve GSE treatments in this study. Linear models with p-value and $R^2$ displayed on the graph.

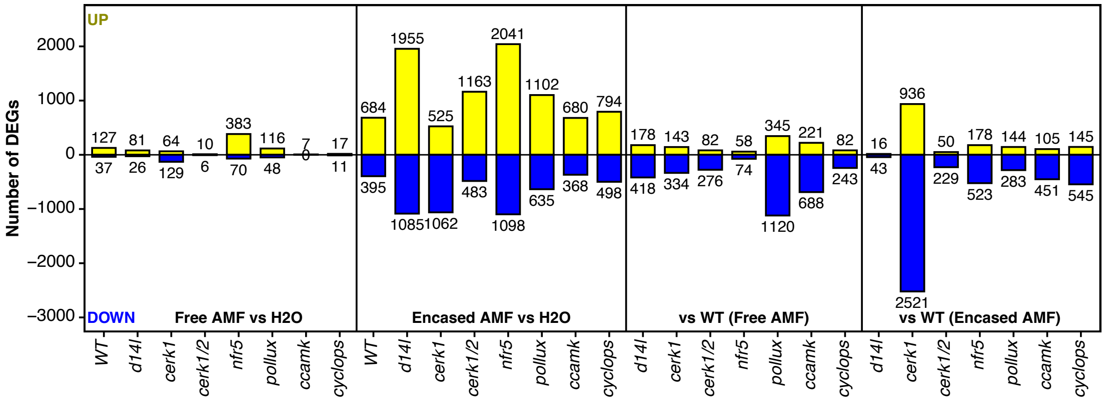


**Fig. S6. Number of differentially expressed genes for AM mutants under encased AMF or free AMF treatments.** Number of differentially expressed genes (DEGs), both up-regulated (positive values, yellow) and down-regulated (negative values, blue), for selected comparisons for AM mutants; in this order: free AMF vs water comparisons, encased AMF vs water, comparisons against wild-type under free AMF treatment, comparisons against wild-type under encased AMF. Numbers over or below bars indicate the precise number of up or down-regulated genes in each comparison.

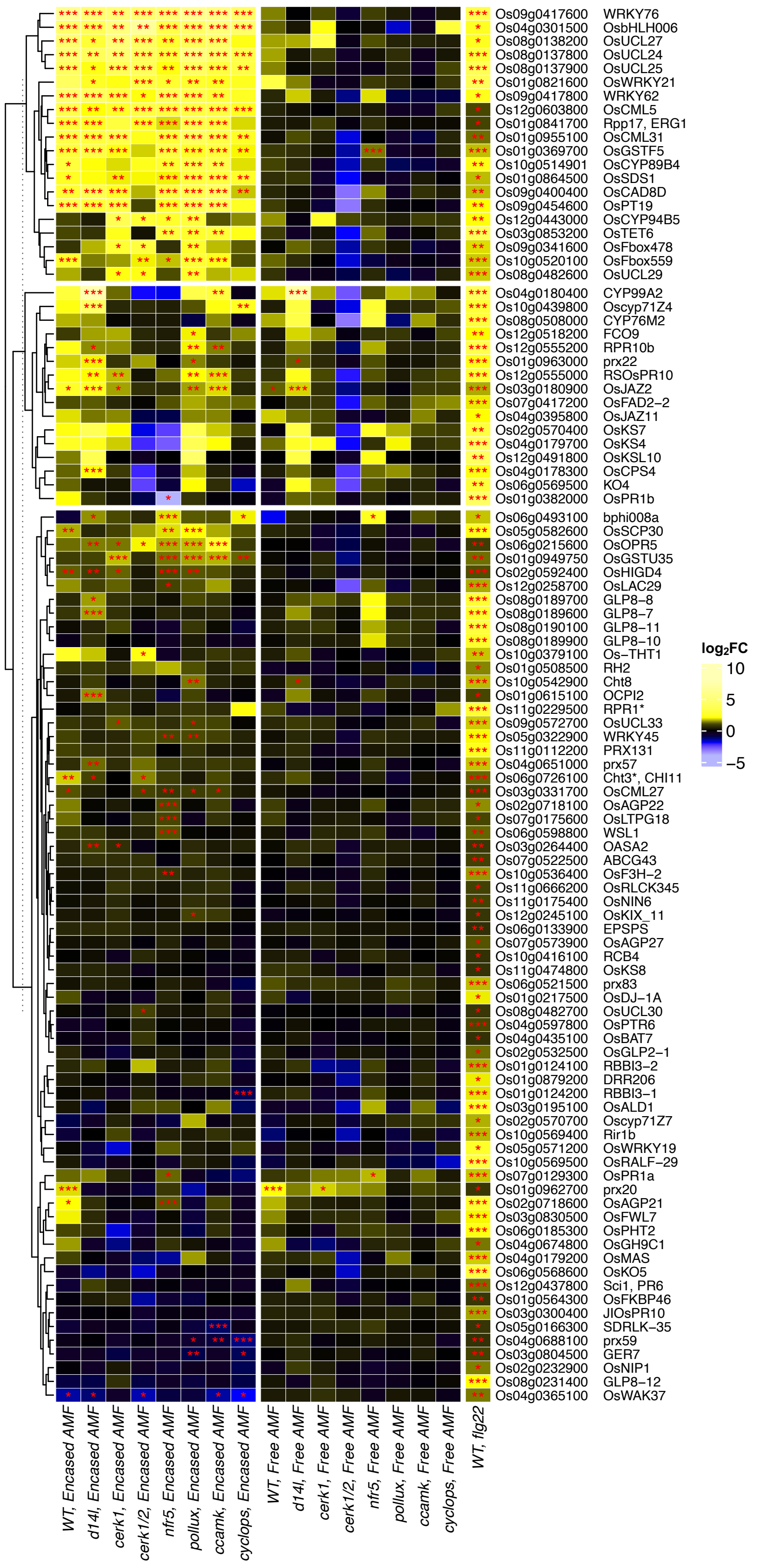


**Fig. S7. PTI signalling is partially activated in AM mutants under encased AMF treatments similarly to wild-type.** Heatmap of $log_2$FC for encased AMF treatments against water for wild-type and AM mutants, as well as flg22 and free AMF treated wild-type against water, for a selection of genes upregulated both under flg22 treatment in this study and *M. oryzae* infection in leaves (Yang et al., 2021 (3)). Significance levels as determined by DEG analysis by DESeq2 shown with asterisks (* for p-value <0.05, ** for < 0.01, *** for < 0.001). Genes were subjected to hierarchical clustering followed by k-means partitioning to subdivide the set in three groups, dendrogram show on the left.

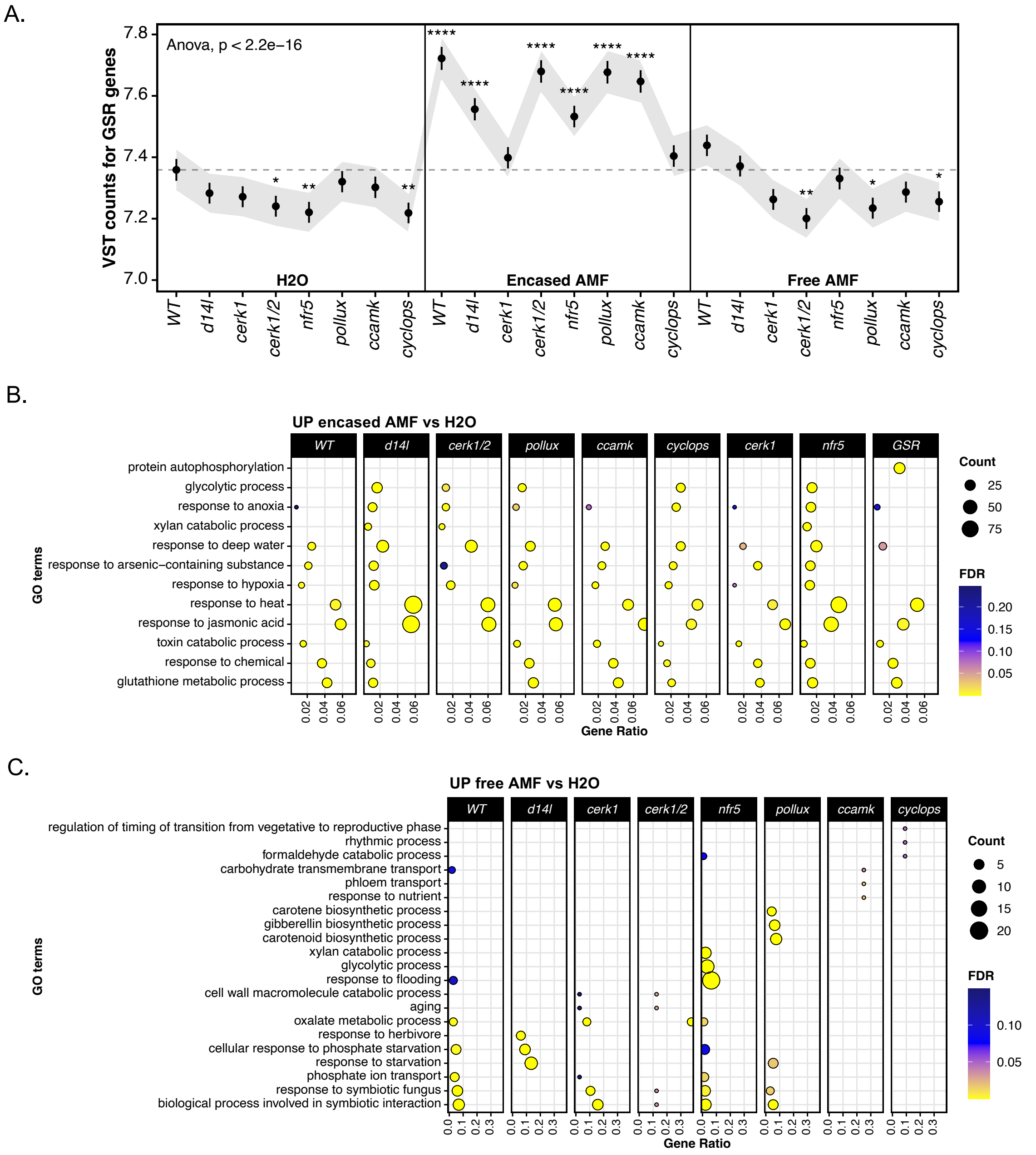


**Fig. S8. AM mutants show a general stress response under encased AMF similar to wild-type, which is not present under free AMF.** (A) Average variance-stabilised count plot for wild-type and mutants exposed to H2O, encased AMF and free AMF for rice orthologues of General Stress Response (GSR) genes (Bjornson et al., 2021 (6)), dots indicate average VST counts, errorbars indicate standard error, shadow indicates normal-based 95% confidence interval around the mean. An ANOVA test was performed followed by pairwise t-test between each condition and wild-type H2O control, significance levels shown with asterisks (* for p-value <0.05, ** for < 0.01, *** for < 0.001 and **** for <0.0001). (B and C) GO term enrichment plot for up-regulated genes in encased AMF (B) and free AMF (C) treatment for all mutants against their respective H2O controls; top three GO terms by gene ratio for each comparison shown, also including wild-type encased AMF induced genes and rice GSR genes as reference (B) or wild-type free AMF (C); colour scale indicates significance as q-value or FDR; dot size proportional to the number of genes in each set included in the GO term; x-axis indicates the ratio of genes up-regulated in each set included in the GO term to the total number of genes annotated with the GO term.

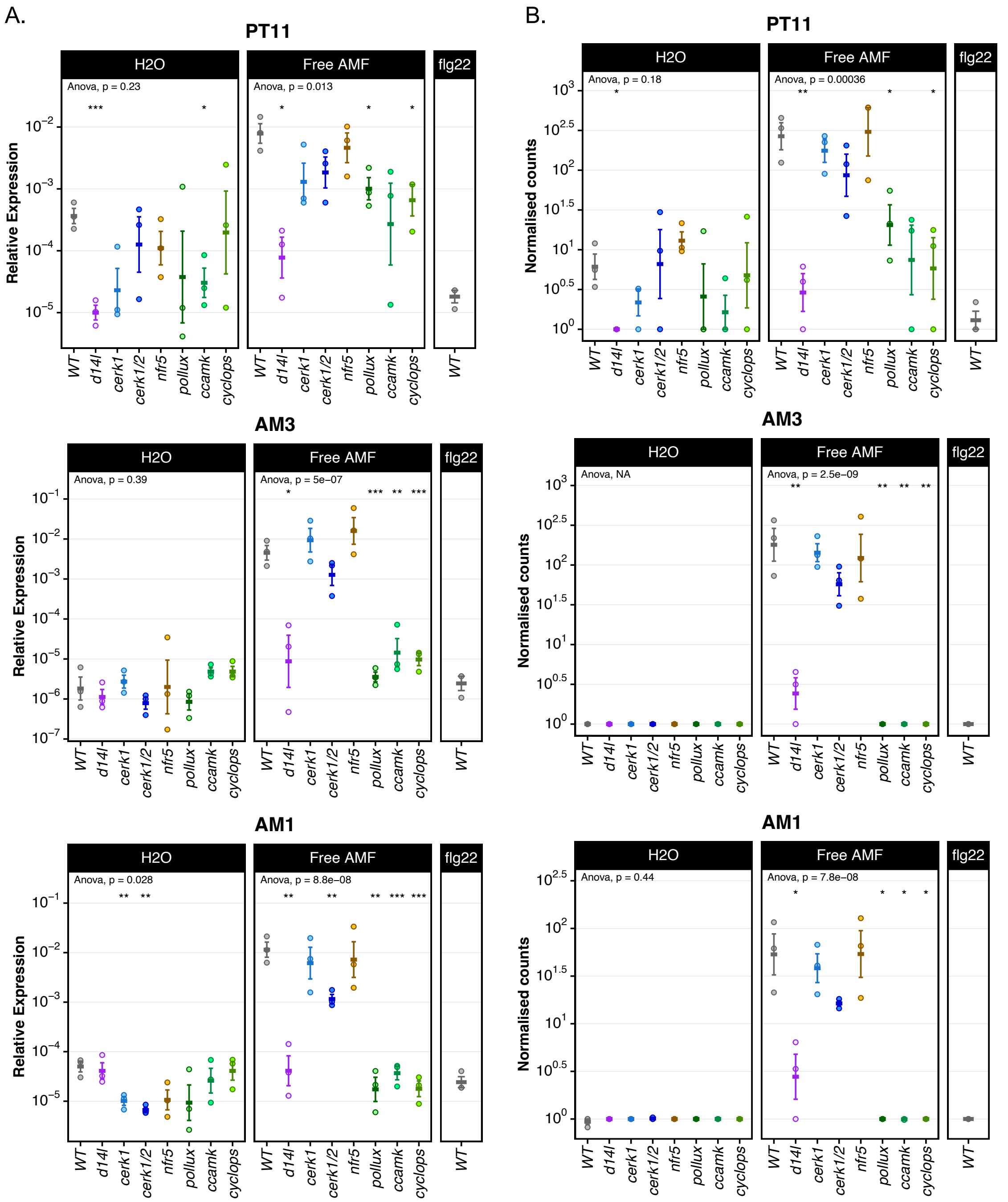


**Fig. S9. Validation of RNA-sequencing results via qRT-PCR.** (A) Relative expression values of *PT11*, *AM1* and *AM3* marker genes as determined by RT-qPCR for same RNA samples used for RNA-sequencing for wild-type and all mutants in H2O and free AMF treatments, and flg22 for the wild-type. (B) DESeq2 normalised counts of *PT11*, *AM1* and *AM3* marker genes as determined by RNA-sequencing for wild-type and all mutants in water and free AMF treatments, and flg22 for the wild-type. Dots indicate individual data biological replicates (average of technical triplicates for qRT-PCR results), crossbar indicates mean, errorbars indicate standard error. An ANOVA test was performed followed by pairwise t-test between each mutant and wild-type for each treatment, significance levels shown with asterisks (* for p-value <0.05, ** for < 0.01, *** for < 0.001 and **** for <0.0001).

**Table S1. Primers used in this study.**

| **Gene** | **Gene ID** | **Primer sequence (5'-3')<br>Forward then reverse** |
|---|---|---|
| For RNA extraction and cDNA synthesis quality controls | | |
| *OsCP2* | *Os02g0121300* | TCCCAGTTCTTCATCTGCAC<br>ACCAAACCATGGGGGATCT |
| *OsGAPDH* | *Os08g0126300* | AGGTTCCTGATTTGAATG<br>CAACTGCACTGGACGGCTTA |
| For RT-qPCR | | |
| *OsACTIN* | *Os03g0718100* | TCTAATTCTTCGGACCCAAGAATG<br>AGCAGGAGGACGGCGATAA |
| *OsCP2* | *Os02g0121300* | GTGGTGTTAGTCTTTTTATGAGTTCGT<br>ACCAAACCATGGGCGATCT |
| *OsPolyubiquitin* | *Os06g0681400* | CATGGAGCTGCTGCTGTTCTAG<br>CAGACAACCATAGCTCCATTGG |
| *OsAM1* | Os04g0134800 | ACCTCGCCAAAATATATGTATGCTATT<br>TTTGCTTGCCACACGTTTTAA |
| *OsAM3* | Os01g0783000 | CTGTTGTTACATCTACGAATAAGGAGAAC<br>CAACTCTGGCCGGCAAGT |
| *OsPT11* | Os01g0657100 | GAGAAGTTCCCTGCTTCAAGC<br>CATATCCCAGATGAGCGTATCATG |
